# *Bifidobacterium pseudocatenulatum* extracellular vesicles promote Ly6G+ granulocyte infiltration to inhibit melanoma tumour progression

**DOI:** 10.64898/2026.06.08.731027

**Authors:** Alicia D. Nicklin, Anne Jordan, Christopher A. Price, Mitchel Rowe, Nilda Ilker, Luke Mitchell, Regis Stentz, Simon R. Carding, Lindsay J. Hall, Stephen D. Robinson

## Abstract

Harnessing the immunomodulatory capacity of commensal bacteria is an emerging avenue in cancer therapy. Bacterial extracellular vesicles (BEVs) provide a non-replicating, nanoscale alternative to live microbes with the potential for safer systemic delivery. Here, we investigated BEVs from a novel Gram-positive strain of *Bifidobacterium pseudocatenulatum* (Bif-210). Intravenous administration of Bif-210 BEVs reduced B16-F10 melanoma growth in C57BL/6J mice. Mechanistically, BEVs increased tumour-infiltrating Ly6G+ granulocytes *in vivo*, increased CD11b+Ly6G+ and ICAM-1+Ly6G+ bone marrow populations, and induced production of the neutrophil-attracting chemokines KC/CXCL1 (mouse) and IL-8 (human). Although Ly6G+ depletion independently inhibited tumour growth, it did not combine additively with BEVs, supporting a model in which Bif-210 BEVs alter Ly6G+ granulocyte function rather than simply expanding a conventional pro-tumour granulocyte pool. BEVs activated TLR2, did not activate TLR4, and upregulated TLR2 on Ly6G+ cells, while proxy assays provided no evidence of NETosis-associated activation. Repeated intravenous BEV administration produced no overt toxicity by tissue histology, body temperature, or body weight monitoring. These findings position B. pseudocatenulatum BEVs as a promising systemic immunotherapy that recruits and re-educates granulocytes via a TLR2-centred pathway to restrain melanoma progression.

**HIGHLIGHTS:**

- Intravenous Bif-210 BEVs reduce established B16-F10 melanoma growth in mice.
- Bif-210 BEVs selectively increase tumour-associated Ly6G+ granulocytes.
- BEV treatment and Ly6G depletion are non-additive, linking BEV activity to granulocyte biology.
- Bif-210 BEVs expand Ly6G+ bone marrow populations and induce granulocyte-recruiting chemokines.
- Bif-210 BEVs engage TLR2 and enhance granulocyte fitness without NETosis-associated activation.

## INTRODUCTION

Members of the genus *Bifidobacterium* are Gram-positive anaerobes that contribute to host health through diverse immune-modifying activities, including effects on epithelial barrier function, microbial metabolism, innate immune activation and adaptive immune education ^1, 2, 3, 4^. Comparative genomics has revealed extensive genetic diversity within the genus, underpinning strain-specific biological activities ^5^ Much of the work to date has focused on *B. bifidum, B. longum, B. breve, B. adolescentis*, and *B. animalis* ^1, 6^. However, less-studied species and novel strains within them represent an underexplored source of host-microbe immune-modulatory biology. Importantly, beneficial effects of *Bifidobacterium* are not limited to live cells: their secreted extracellular vesicles (EVs) can also mediate host-facing biological activities ^7, 8^.

Bacterial extracellular vesicles (BEVs) are nanosized, membrane-bound particles released by Gram-negative and Gram-positive bacteria ^9^. While early studies focused on Gram-negative BEVs, increasing attention has turned to Gram-positive BEVs ^10^. This shift is clinically relevant: Gram-negative BEVs may overstimulate the immune system through virulence factors such as LPS and leukotoxins ^11^, whereas commensal Gram-positive BEVs could offer a more favourable safety profile ^12^. BEV composition is shaped by bacterial species, strain, growth phase and culture conditions, and these differences critically influence host responses ^13, 14, 15^.

Immunomodulation is among the most prominent BEV-host interactions, but immunomodulation is not a single response: BEVs can influence cytokine production, pattern-recognition receptor signalling, antigen presentation, cell recruitment, myeloid cell activation and adaptive immune priming depending on the vesicle source and recipient cell context ^16, 17^. Detailed characterisation of *Bifidobacterium* BEVs remains limited. EVs from *B. longum* increased IL-6 via TLR2 activation in macrophage-like RAW264.7 cells ^18^, while vesicles from distinct *B. longum* strains have been reported to exert anti-inflammatory and tissue-distribution properties that depend on the vesicle source and experimental context ^19, 20, 21^. Given that *Bifidobacterium* is the dominant genus in the infant gut, a period when immune priming is especially active, its EVs may have particular immunomodulatory potential ^22^.

The concept of using microbes or microbial products in cancer therapy is well established. *Mycobacterium bovis bacille Calmette-Guerin* (BCG) remains a mainstay treatment for non-muscle-invasive bladder cancer ^23^, and BCG-derived EVs can replicate some immune-stimulating effects of live BCG ^24^. Similarly, a patient’s gut microbiota composition, including *Bifidobacterium*, has been linked to clinical responses to immune checkpoint blockade in melanoma ^25, 26^. In mouse models, oral *Bifidobacterium* reduced melanoma burden, potentially through priming of cytotoxic CD8+ T cells ^27^, and *B. longum* administration suppressed colon carcinogenesis ^28, 29^. More recent studies directly implicate *Bifidobacterium*-derived EVs in cancer biology, including protection against hepatocellular carcinoma-associated pathways ^30^, modulation of anti-PD-1 therapy in lung cancer ^31^, and inhibition of colorectal cancer by *B. breve* EVs containing formate acetyltransferase ^32^. These studies support the broader antitumour potential of *Bifidobacterium* and its vesicle-mediated effects, while leaving the activity of *B. pseudocatenulatum* BEVs largely unexplored.

A major unresolved question is which immune compartments can be productively redirected by microbial vesicles. Ly6G+ granulocytes, often referred to as neutrophils in murine tumour models, are especially relevant because their functions in cancer are highly context-dependent. The N1/N2 tumour-associated neutrophil (TAN) framework was established by studies showing that TGF-beta modulation can generate CD11b+Ly6G+ neutrophils with enhanced antitumour activity ^33^, and transcriptomic analyses have since shown that pro- and antitumour TAN states are distinct and dynamic ^34^. This plasticity creates an opportunity for microbial products to recruit granulocytes while also shifting their functional state.

Here, we isolated BEVs from a novel *B. pseudocatenulatum* strain, Bif-210, and tested their immunotherapeutic effects in a murine melanoma model. We first asked whether systemic Bif-210 BEV administration could inhibit established tumour growth, then used immune profiling and mechanistic assays to identify the immune compartment most strongly associated with the response. Rather than presupposing a neutrophil-centred mechanism, our data identified Ly6G+ granulocytes as the dominant population altered by BEV treatment and led us to test whether Bif-210 BEVs reprogramme granulocyte biology through a TLR2-centred pathway.

## RESULTS

### Intravenous Bif-210 BEVs suppress established melanoma growth

To test whether Bif-210 BEVs influence established tumour growth, C57BL/6J mice bearing subcutaneous B16-F10 melanomas received intravenous injections of BEVs or control (PBS) beginning on day 6, when tumours first became palpable. Injections were repeated on days 9 and 12 (Fig. 1A). This design was intended to reflect a therapeutically relevant setting in which treatment begins after tumour establishment, while intravenous delivery maximised systemic exposure to circulating immune cells and tumour tissue.

**Figure 1.**
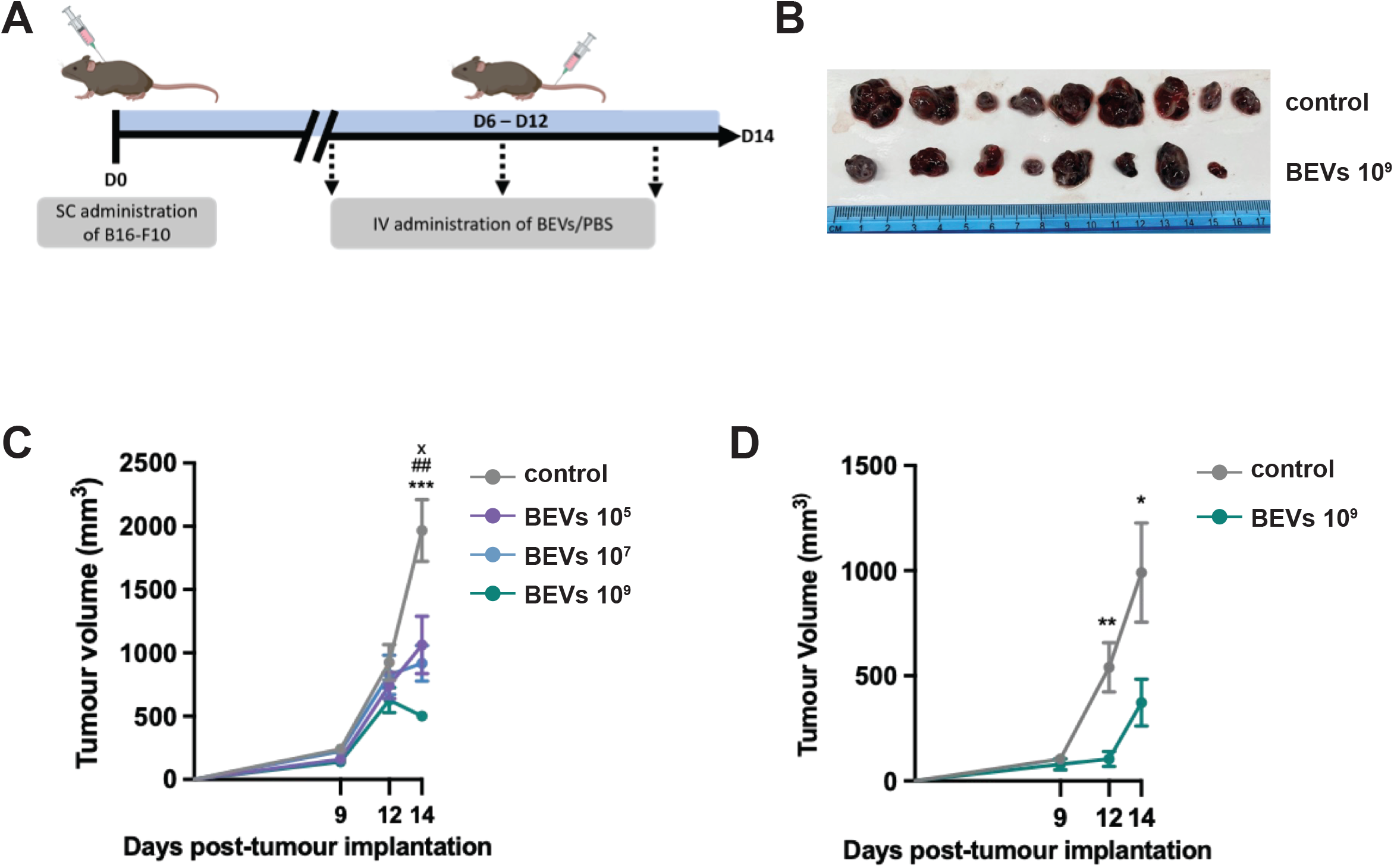
Bif-210 BEVs suppress established melanoma growth. (A) Experimental timeline. B16-F10 cells were injected s.c. on day 0. BEVs or PBS were administered i.v. on days 6, 9 and 12. Mice were culled on day 14. (B) Representative resected tumours from PBS and BEV-treated (10^9^ particles/mouse) groups. (C) Dose-response study showing tumour volume in PBS vs BEVs (10^5^, 10^7^, 10^9^ particles/mouse) at days 9, 12 and at resection (day 14). (D) Fixed-dose (10^9^ particles/mouse) efficacy over time (days 9, 12) and at resection (day 14). Tumour volume = width^2^ × length × 0.52. Statistics: (C) multiple unpaired t-tests; (D) two-way ANOVA with Dunnett’s post hoc (vs PBS). Significance vs PBS is denoted by * or x (p<0.05), ** or ## (p<0.01), *** (p<0.005).

Mice treated with Bif-210 BEVs exhibited significantly smaller tumours than PBS controls, with reductions evident from day 12 (Fig. 1B-D). A dose-response experiment using 10^5^, 10^7^ and 10^9^ BEVs per mouse showed that each dose suppressed tumour growth, with 10^9^ particles producing the most pronounced tumour suppression (Fig. 1C). This dose was therefore selected for subsequent mechanistic experiments.

Because bacterial products delivered systemically can induce inflammatory toxicity, we also assessed repeated intravenous dosing in non-tumour-bearing mice. Using the same 10^9^-particle dosing schedule, Bif-210 BEVs caused no overt tissue pathology in liver, spleen, kidney or lung (Fig. S1A), and did not alter body temperature (Fig. S1B) or body weight (Fig. S1C) over the treatment period. These data support the tolerability of systemic Bif-210 BEV delivery at the therapeutic dose used in this study.

### Bif-210 BEVs converge on tumour-associated Ly6G+ granulocyte biology

Having established antitumour activity and initial tolerability, we next profiled tumour immune infiltrates to determine how Bif-210 BEVs reshape the tumour microenvironment. Flow cytometric immunophenotyping revealed a significant increase in Ly6G+ granulocytes in tumours from BEV-treated mice compared with PBS controls across three independent experiments (Fig. 2A-B). By contrast, other major myeloid and lymphoid subsets, including B-cell, T-cell, and dendritic-cell populations, were largely unchanged (Fig. 2C). These data indicate that Bif-210 BEVs primarily alter the tumour immune microenvironment through the granulocyte compartment.

**Figure 2.**
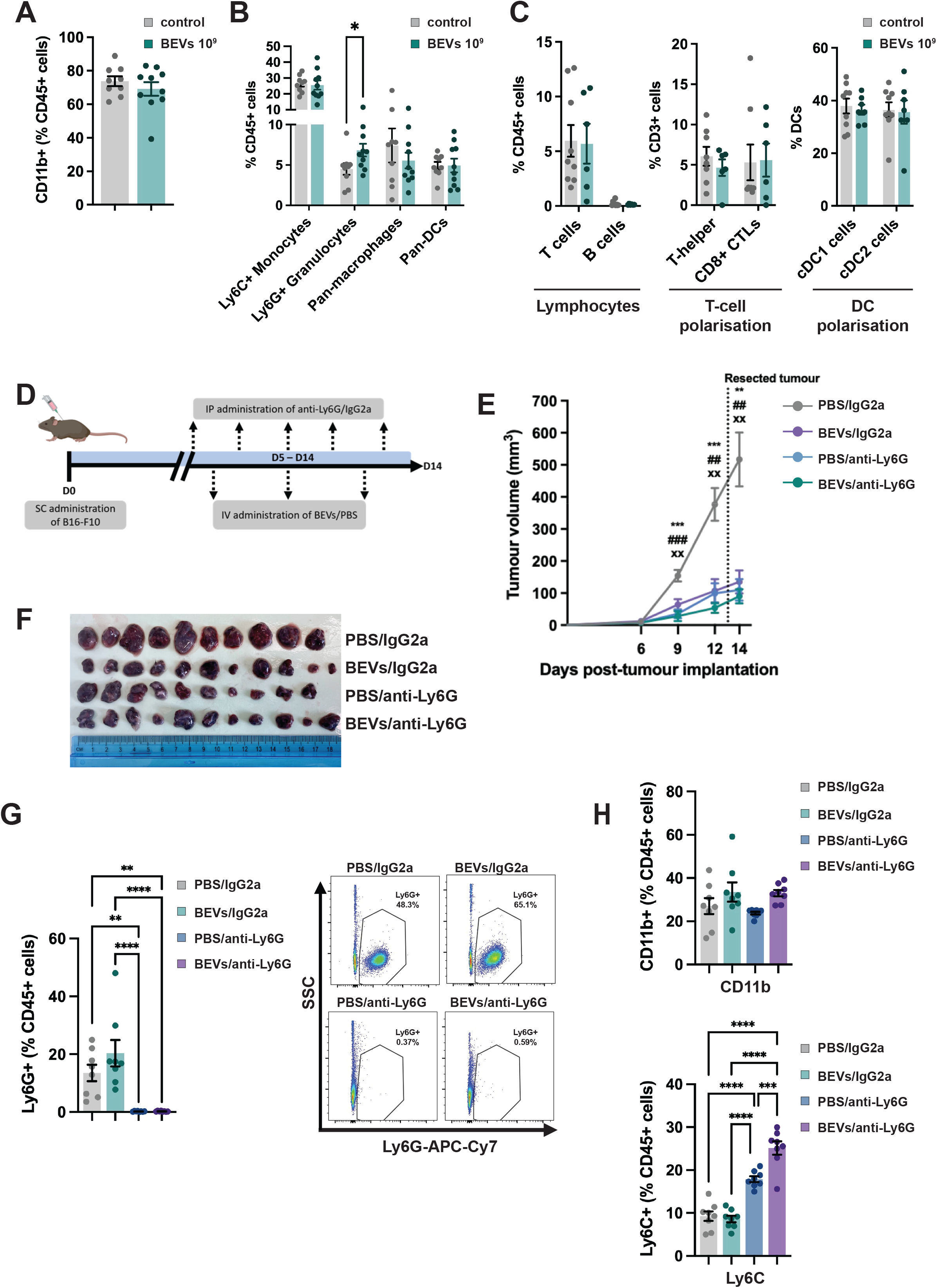
Bif-210 BEVs converge on tumour-associated Ly6G+ granulocyte biology. (A) Myeloid populations within tumours expressed as % of CD45+ cells (PBS vs BEVs, 10^9^ particles/mouse). (B) Ly6G+ granulocytes pooled across three independent experiments. Each point represents an individual tumour and is expressed relative to the matched PBS-treated control mean; representative dot plots illustrate the Ly6G+ shift after BEV treatment. (C) Other tumour immune populations, including lymphocyte populations, T-cell polarisation and dendritic-cell polarisation, showing no broad remodelling of major non-granulocyte compartments. (D) Ly6G depletion experimental design. B16-F10 cells were injected s.c. on day 0. Anti-Ly6G (clone 1A8, 200 µg i.p.) or IgG2a isotype control was started on day 5 and administered every 2 days. BEVs or PBS were given i.v. on days 6, 9 and 12. Mice were culled on day 14. (E) Tumour volumes for PBS/IgG2a, BEVs/IgG2a, PBS/anti-Ly6G and BEVs/anti-Ly6G groups. Two-way ANOVA with Tukey’s multiple comparisons. Symbols in the panel denote comparisons vs PBS/IgG2a. (F) Representative resected tumours from all four depletion/treatment groups. (G) Blood Ly6G+ (% CD45+) confirms efficient depletion (<1% in anti-Ly6G groups); representative gating shown. (H) Blood myeloid compartment, including CD11b+ myeloid cells and Ly6C+ monocytes (% of CD45+), following BEV and/or anti-Ly6G treatment. Bars = mean ± SEM; statistical tests as indicated in Methods and original panels.

The increase in tumour-associated Ly6G+ cells created an apparent paradox, as Ly6G+ granulocytes can either promote or restrict tumour growth depending on context. To define how this population contributed to melanoma progression in our model, Ly6G+ cells were depleted *in vivo* using an anti-Ly6G antibody (clone 1A8), with BEVs or PBS administered on the same schedule as above (Fig. 2D). Ly6G+ depletion alone significantly inhibited tumour growth, and BEV monotherapy produced a comparable reduction in tumour burden (Fig. 2E-F). Importantly, combining BEVs with Ly6G depletion did not further reduce tumour burden beyond either intervention alone (Fig. 2E). Peripheral blood flow cytometry confirmed efficient depletion of Ly6G+ cells (Fig. 2G). Additional circulating myeloid changes observed after combined BEV and anti-Ly6G treatment are shown in Fig. 2H. Together, these findings suggest that Bif-210 BEVs converge on Ly6G+ granulocyte biology rather than acting through a completely independent pathway, supporting a model in which BEVs alter the functional state of Ly6G+ cells.

### Bif-210 BEVs expand Ly6G+ bone marrow populations and induce granulocyte-recruiting chemokines

Because circulating granulocytes arise from the bone marrow, we next investigated whether Bif-210 BEVs act directly on bone marrow immune populations. *Ex vivo* exposure of bone marrow cells to BEVs for five hours markedly increased the proportion of CD11b+Ly6G+ cells within the CD45+ population, from a small baseline population to a majority of CD45+ cells (Fig. 3A). BEV stimulation also increased ICAM-1 expression on Ly6G+ cells (Fig. 3B), consistent with acquisition of an ICAM-1-high, N1-like phenotype rather than simple expansion of suppressive granulocytes ^33, 34, 35^. In line with these *ex vivo* observations, bone marrow analysis from non-tumour-bearing and tumour-bearing mice also supported BEV-driven changes in CD11b+ and Ly6G+ populations within the CD45+ compartment (Fig. 3C-D).

**Figure 3.**
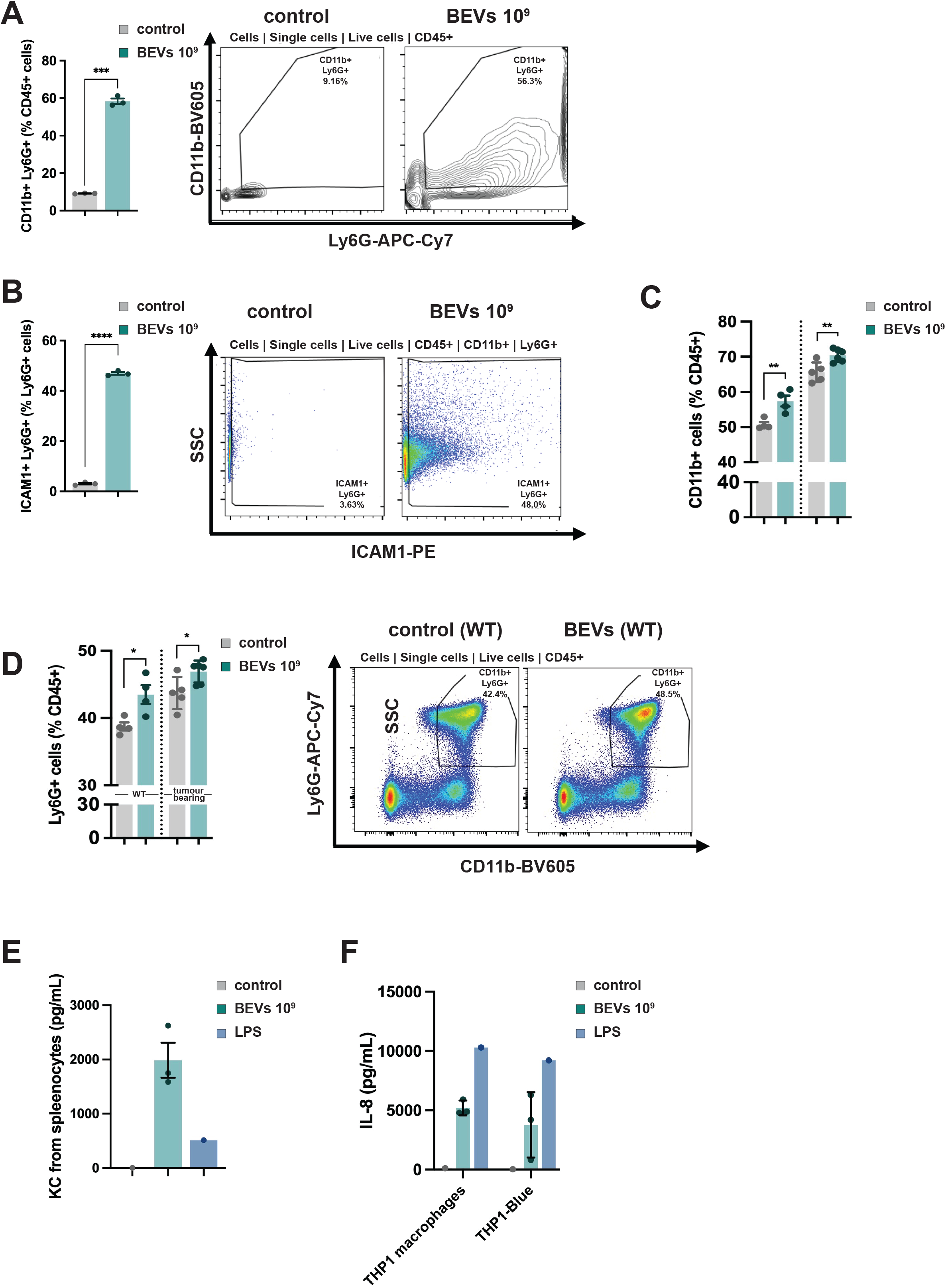
Bif-210 BEVs expand Ly6G+ bone marrow populations and induce granulocyte-recruiting chemokines. (A) Bone marrow cells were incubated ex vivo with PBS or BEVs (10^9^ particles/well) for 5 h. CD11b+Ly6G+ cells among CD45+ cells were quantified by flow cytometry; representative contour plots shown. (B) ICAM-1 expression on Ly6G+ cells after BEV exposure; representative plots and quantification shown. (C-D) *In vivo* bone marrow CD11b+ and Ly6G+ populations among CD45+ cells in non-tumour-bearing and tumour-bearing mice following PBS or BEV administration. (E) KC/CXCL1 levels in mouse splenocyte supernatants following PBS, BEV or LPS stimulation. (F) IL-8 levels in THP-1 macrophage and THP1-Blue supernatants following PBS, BEV or LPS stimulation. Statistics: (A-B) paired two-tailed t-tests; other statistical tests as indicated in Methods. Bars = mean ± SEM.

To determine how increased granulocyte infiltration into tumours might be driven, we measured neutrophil-recruiting chemokines following BEV stimulation of immune cells. Murine splenocytes secreted high levels of keratinocyte-derived cytokine (KC/CXCL1) after BEV exposure, exceeding levels induced by LPS under these conditions (Fig. 3E). Similarly, human THP-1 macrophages and THP1-Blue monocytes secreted elevated IL-8 following BEV stimulation (Fig. 3F). These data suggest that Bif-210 BEVs influence both granulocyte generation/phenotype in the bone marrow and chemokine-mediated recruitment signals that could support tumour infiltration.

### Bif-210 BEVs engage TLR2 and enhance granulocyte fitness without NETosis-associated activation

We next examined the innate receptor pathway through which Bif-210 BEVs activate immune cells. Bone marrow cultures exposed to BEVs showed increased TLR2 expression on Ly6G+ cells (Fig. 4A). Consistent with this, HEK-Blue reporter assays demonstrated that Bif-210 BEVs activated TLR2 at the treatment-relevant dose, whereas BEVs produced no detectable TLR4 activation under matched conditions (Fig. 4B-C). These findings support a TLR2-centred pathway, consistent with the Gram-positive composition of Bif-210 BEVs. The absence of detectable TLR4 activation should not be interpreted as implying that all TLR4 signalling is detrimental, because distinct LPS structures can produce different immunological outcomes, including immunotherapy-enhancing effects of hexa-acylated LPS ^36^. Rather, in this system, Bif-210 BEVs preferentially engage TLR2 under the conditions tested.

**Figure 4.**
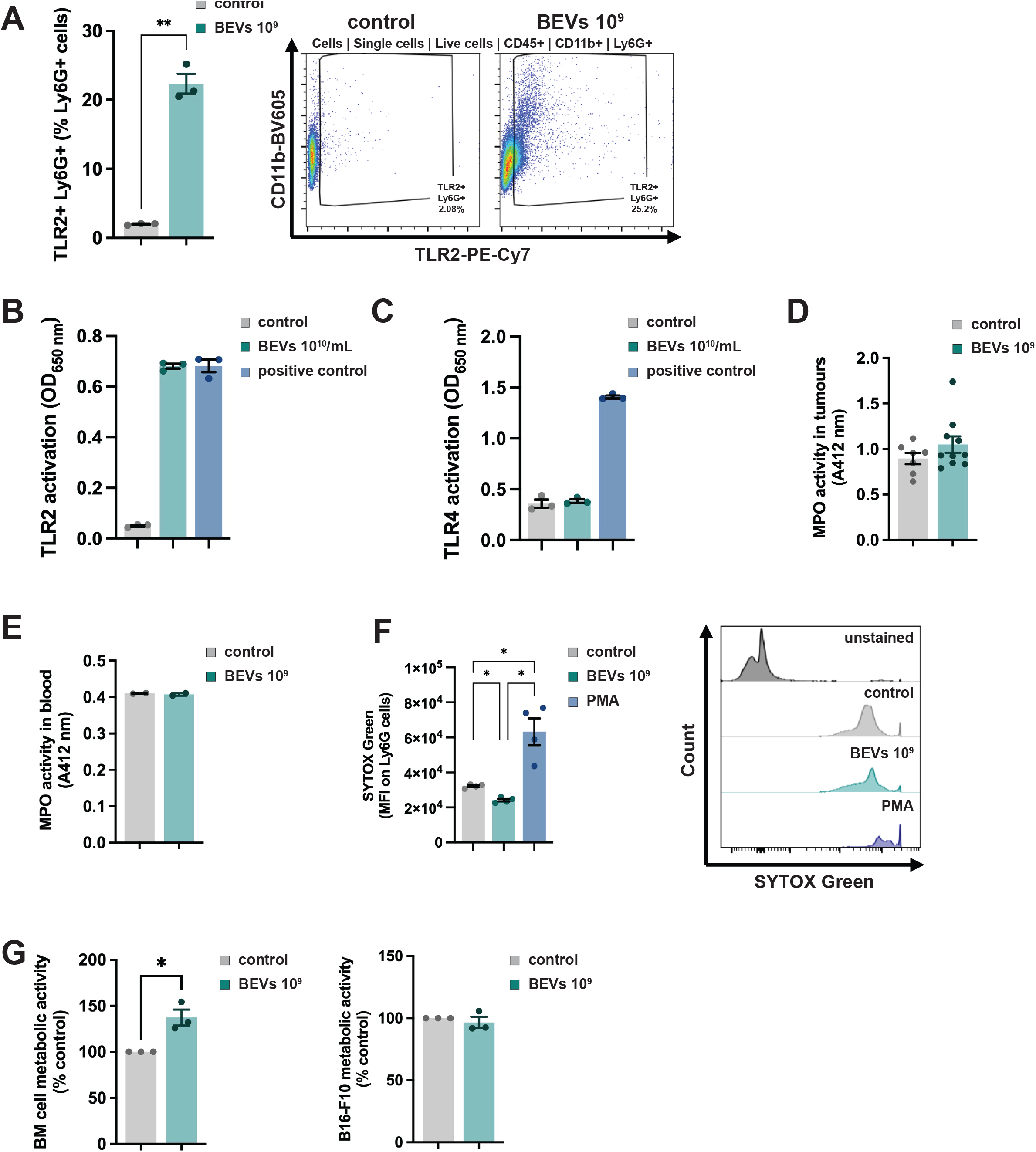
Bif-210 BEVs engage TLR2 and enhance granulocyte fitness without NETosis-associated activation. (A) TLR2 expression on bone marrow-derived Ly6G+ cells after 5 h incubation with PBS or BEVs; representative plots and quantification shown. (B) HEK-Blue hTLR2 reporter assay showing BEV-driven TLR2 activation at the treatment-relevant dose, with LTA-BS as a positive control. The full dose-response is shown in Fig. S2. (C) HEK-Blue hTLR4 reporter assay showing no BEV-driven TLR4 activation under matched conditions; LPS served as positive control. (D) MPO activity in B16-F10 tumour lysates from PBS- and BEV-treated mice. (E) MPO activity in mouse whole blood incubated ex vivo with PBS or BEVs (10^9^ particles). Absorbance at 412 nm is proportional to MPO activity. (F) Extracellular DNA release from murine bone marrow cells assessed by SYTOX Green flow cytometry after PBS, BEV or PMA treatment. PMA served as a positive control for NETosis-associated extracellular DNA release. (G) MTS assay assessing metabolic activity/viability of murine bone marrow cells and B16-F10 tumour cells following PBS or BEV exposure. Values are expressed relative to matched PBS controls. Statistics as indicated in Methods and original panels.

Because TLR2 activation can in some contexts promote neutrophil extracellular trap formation, we next assessed NETosis-associated readouts. Myeloperoxidase (MPO) activity, used as a surrogate of degranulation or NET-associated granule release, was not increased in tumour homogenates from BEV-treated mice or in whole blood incubated ex vivo with BEVs (Fig. 4D-E). BEVs also did not increase SYTOX Green-positive extracellular DNA release from bone marrow cultures, whereas PMA induced robust extracellular DNA release (Fig. 4F; ^37, 38^. Finally, MTS assays showed that BEV-treated bone marrow cells had higher metabolic activity than PBS controls, whereas B16-F10 tumour cells showed no direct change in metabolic activity/viability following BEV exposure (Fig. 4G). These proxy assays provide no evidence that Bif-210 BEVs induce a NETosis-associated granulocyte state; instead, they support enhanced bone marrow immune-cell viability and function.

## DISCUSSION

This study provides the first evidence that extracellular vesicles derived from a novel *B. pseudocatenulatum* strain (Bif-210) suppress melanoma growth in a preclinical mouse model. Systemic Bif-210 BEV administration reduced tumour burden in the B16-F10 model without directly reducing B16-F10 cell metabolic activity/viability, supporting an immune-mediated mechanism rather than direct tumour-cell cytotoxicity.

A key biological theme emerging from these data is granulocyte plasticity. Bif-210 BEVs increased tumour-associated Ly6G+ granulocytes, yet Ly6G depletion independently reduced tumour growth. This could appear contradictory if Ly6G+ cells are treated as a single functional entity. However, tumour-associated neutrophils (TANs) and granulocytic myeloid populations can adopt pro- or antitumour states depending on local signals, as shown by N1/N2 polarisation studies and transcriptomic analyses of TAN states ^33, 34, 39^. The absence of additive benefit when BEVs were combined with antibody-mediated Ly6G cell depletion suggests that these interventions converge on granulocyte population dynamics and function. A cautious interpretation is that Bif-210 BEVs alter the functional properties of Ly6G+ cells rather than simply increasing the abundance of a conventional pro-tumour granulocyte population.

Several observations support this model. First, BEVs increased CD11b+Ly6G+ bone marrow populations and upregulated ICAM-1 on Ly6G+ cells, consistent with an activated or N1-like phenotype ^35^. Second, BEVs induced KC/CXCL1 and IL-8, suggesting a chemokine axis capable of recruiting granulocytes from bone marrow and blood into tumours. Third, BEVs activated TLR2 but not TLR4 and increased TLR2 expression on Ly6G+ cells. Collectively, these findings are consistent with the Gram-positive composition of Bif-210 BEVs and suggest a TLR2-mediated activation pathway ^40, 41^. At the same time, the lack of TLR4-associated activation should be interpreted specifically for Bif-210 BEVs rather than as a general statement about Gramnegative BEVs or LPS biology, because structurally distinct LPS species can elicit divergent effects on tumour immunity and checkpoint blockade responses ^36^. Although Bif-210 BEVs engaged TLR2 under the conditions tested, contributions from additional pattern-recognition receptors, including other TLRs or NLRs, cannot be excluded.

Importantly, the BEV-induced Ly6G+ response was not associated with NETosis proxy readouts. NET formation can promote tumour growth, metastasis and inflammatory tissue damage ^42^. Bif-210 BEVs did not increase MPO activity or extracellular DNA release and instead enhanced bone marrow cell metabolic activity. These findings help distinguish the BEV response from generic neutrophil activation and support a more selective granulocyte-reprogramming mechanism.

The tolerability data strengthen the translational rationale for systemic Bif-210 BEV administration. No overt histological abnormalities were observed in major organs, and body temperature and weight remained stable following repeated 10^9^-particle dosing. These data are encouraging, but should be interpreted as an initial safety screen rather than a full toxicology package. Future studies should include serum inflammatory cytokines, clinical chemistry, longer follow-up and dose-escalation analyses, especially if intravenous delivery is pursued.

From a translational perspective, BEVs offer advantages over live bacteria. As non-replicating particles, they reduce risks associated with infection, persistence, environmental contamination and horizontal gene transfer ^43^. However, their therapeutic development will depend on robust batch control. BEV composition can vary with bacterial growth phase and culture conditions ^13, 15^, and current EV reporting standards emphasise the need for complementary particle, biophysical, biochemical, immunological and purity measurements ^17^. For Bif-210 BEVs, future optimisation should include particle-size distribution, protein/lipid content, endotoxin testing, sterility and cargo fingerprinting. To avoid pre-empting detailed BEV characterisation data being developed in parallel, the present manuscript reports minimal BEV characterisation required for interpretation of the melanoma experiments; these data are further developed and discussed in a broader context in a separate manuscript (A. Jordan et al., manuscript in preparation).

Clinically, Bif-210 BEVs may be most attractive as innate immune-modulating adjuncts to existing immunotherapies. Microbiota-associated enhancement of checkpoint blockade has been linked to improved antitumour immunity in melanoma ^25, 26, 27^ and BEV-mediated granulocyte reprogramming could complement adaptive immune activation. Further insight into the mechanism of action of Bif-210 BEVs will require more detailed analyses of BEV-conditioned tumour-associated Ly6G+ cells, including their cytotoxicity, transcriptomic profile, cytokine production and ability to cooperate with CD8+ T cells. This will be important for converting the current model from a compelling association into a directly demonstrated effector mechanism.

In conclusion, Bif-210 BEVs represent a promising platform for microbial vesicle-based cancer immunotherapy. By recruiting Ly6G+ granulocytes, promoting TLR2-mediated myeloid activation and avoiding NETosis-associated readouts or overt systemic toxicity, these vesicles provide a route to exploit innate immune plasticity for tumour control.

## LIMITATIONS OF THE STUDY

Bone marrow Ly6G+ cells provide a practical and reproducible system for assessing granulocyte responses because they can be isolated in sufficient numbers and handled under controlled *ex vivo* conditions. However, tumour-associated Ly6G+ cells are typically less abundant, short-lived and sensitive to tissue dissociation, and future studies should directly profile tumour-infiltrating Ly6G+ populations where sufficient viable cells can be obtained. Direct cytotoxicity assays, transcriptomic profiling and functional re-stimulation analyses would help define whether BEV-conditioned tumour granulocytes acquire a stable antitumour state.

TLR2 activation was demonstrated using reporter assays and increased TLR2 expression on Ly6G+ cells, but the *in vivo* requirement for TLR2 signalling remains to be established. Future studies should test whether interference with TLR2 signalling alters BEV-driven granulocyte activation and tumour control.

Finally, BEV composition and cargo were not molecularly resolved in the current study. Defining the TLR2 ligands and the proteins, lipoproteins, polysaccharides or nucleic acids responsible for Ly6G+ polarisation will be essential for quality control, reproducibility and therapeutic development.

## MATERIALS AND METHODS

### Bacterial strain and culture conditions

*B. pseudocatenulatum* Bif-210 was isolated previously in-house by the laboratory of Prof. Lindsay Hall (from a 159 day old human infant, delivered vaginally). The strain was cultured anaerobically at 37°C in vegan reinforced clostridial medium (RCM) containing yeast extract (13.0 g/L, Thermo), soymeal-based peptone (10.0 g/L, Sigma), soluble starch from potato (1.0 g/L, Sigma), alpha-D-glucose (5.0 g/L, Sigma), cysteine hydrochloride from non-animal source (0.5 g/L, Sigma), sodium chloride (5.0 g/L, Sigma) and sodium acetate (3.0 g/L, Sigma), supplemented with L-cysteine (50 mg/L, Sigma).

### BEV isolation, purification and storage

Bif-210 was inoculated at 5% (v/v) into 500 mL filtered and oxygen-reduced vegan RCM and grown for 20 h at 37°C under humidified anaerobic conditions. Cultures were centrifuged (7,000 g, 1 h, 4°C) and supernatants filtered through 0.22 µm PES bottle-top filters (Sartorius). Supernatants were concentrated using counterflow ultrafiltration through a Vivaflow 50R cassette (100 kDa MWCO; Sartorius) connected to a peristaltic pump. The system was washed with 500 mL PBS and the filtrate was collected and centrifuged. BEVs were further purified by size-exclusion chromatography using qEVoriginal 35 nm columns (Izon) according to the manufacturer’s instructions. After final sterile filtration (0.22 µm PES), aliquots were stored at -80°C. BEV isolation and reporting were guided by published bacterial EV isolation approaches and current EV reporting guidance ^17, 44^.

BEV particle concentration was determined by nanoparticle tracking analysis (NTA) using a ZetaView instrument with software version 8.05.12SP1 (Particle Metrix), as previously described for bacterial extracellular vesicles ^15^. Samples were analysed using a 2-cycle, 11-position, high-frame-rate analysis at 25°C, with fixed camera-control settings (sensitivity 80, frame rate 30, shutter 100) and post-acquisition parameters (brightness 20, maximum area 2000, minimum area 5, trace length 30, 5 nm/class, 64 classes/decade). Dose is reported as particles per mouse or particles per mL. A concise summary of BEV size, concentration and particle-to-protein ratio is provided in Table 1. These BEV characterisation data are further developed and discussed in a broader context in a separate manuscript (A. Jordan et al., manuscript in preparation).

**Table 1.** Minimal characterisation of Bif-210 BEV preparations used in this study.

|  |  |
| --- | --- |
| Parameter | Mean ± SEM |
| Particle concentration ($\times 10^{11}$ particles/mL) | 3.54 ± 0.84 |
| Protein concentration (μg/mL) | 76.7 ± 11.2 |
| Protein per particle ($\times 10^{-10}$ μg/particle) | 2.44 ± 0.42 |
| Particle-to-protein ratio ($\times 10^9$ particles/μg protein) | 4.49 ± 0.77 |
| Weighted mean particle diameter (nm) | 142.9 ± 3.0 |
| Modal particle diameter (nm) | 105 ± 5 |
| Values are shown as mean ± SEM from four independent Bif-210 BEV preparations. Particle concentration and size distribution were determined by nanoparticle tracking analysis. Protein concentration was determined by protein assay. Weighted mean particle diameter was calculated from binned NTA size-distribution data. Modal particle diameter represents the most abundant size bin within each preparation. Broader BEV characterisation data are further developed and discussed in a separate manuscript (A. Jordan et al., manuscript in preparation). | |

### Cell culture

B16-F10 murine melanoma cells (ATCC CRL-6475) were cultured at 37°C/5% CO_2_ in high-glucose DMEM (Thermo Fisher) supplemented with 10% foetal bovine serum (FBS; Hyclone, Thermo Fisher) and 100 units/mL penicillin/streptomycin (P/S; Thermo Fisher). Cells were grown in tissue-culture-treated flasks precoated with 0.1% porcine gelatin (Sigma) and subcultured at approximately 70% confluency.

THP-1 monocytes (ATCC, TIB-202), THP1-Blue NF-kB reporter monocytes and HEK-Blue hTLR2/hTLR4 reporter cells (InvivoGen) were cultured at 37°C/5% CO_2_ in RPMI-1640 (Gibco) containing L-glutamine (2 mM) and HEPES (25 mM), supplemented with heat-inactivated FBS. Reporter lines were maintained with Normocin (100 µg/mL; InvivoGen) and appropriate HEK-Blue/blasticidin selection according to the manufacturer’s instructions. THP-1 cells were differentiated into macrophage-like cells using PMA (10 ng/mL; Sigma) for 24 h.

Murine splenocytes were isolated from C57BL/6J spleens by mechanical homogenisation and passage through 70 µm sterile cell strainers. Cells were washed in sterile PBS and cultured in RPMI-1640 supplemented with 10% FBS, 55 µM beta-mercaptoethanol (Fisher Scientific), 1% P/S, 1 mM sodium pyruvate (Gibco) and GM-CSF (80 ng/mL; Fisher Scientific) at 37°C/5% CO_2_.

### Animals

All animal work was approved by the UK Home Office and performed under project licence PP8873233. Specific-pathogen-free C57BL/6J mice, aged 8-12 weeks, were maintained at the University of East Anglia Disease Modelling Unit (Norwich, UK) with *ad libitum* access to food and water and appropriate environmental enrichment. Both sexes were included in these studies.

For tumour-growth experiments, the experimental unit was the individual mouse. The primary outcome measure was tumour volume. Secondary outcome measures included tumour immune-cell populations, blood myeloid populations, bone marrow Ly6G+ populations, TLR reporter activation, NETosis-associated readouts and safety/tolerability measures. Experimental groups are described in the corresponding figure legends and included PBS controls, Bif-210 BEV-treated mice, isotype control-treated mice and anti-Ly6G-treated mice where appropriate.

Sample sizes were based on previous experience with subcutaneous B16-F10 tumour experiments and flow-cytometric immune profiling, using the minimum number of animals required to detect biologically meaningful differences while respecting ethical reduction of animal use. No formal *a priori* sample size calculation was performed. Exact n values for each experiment and analysis are reported in the figures/figure legends.

Animals were included if they were within the specified age range, had successful tumour-cell implantation, and remained clinically suitable for study until the planned endpoint. Samples were excluded from individual analyses only where technical failure prevented reliable quantification, such as failed tissue recovery, poor flow-cytometry acquisition, insufficient cell yield or unusable histology/imaging quality. No animals or samples were excluded on the basis of the observed tumour phenotype unless prespecified humane endpoints were reached. Exact n values after any exclusions are reported in the relevant figures/figure legends.

Mice were randomly allocated to treatment groups to balance tumour establishment and experimental processing order where possible. Treatment administration could not always be blinded because BEV/PBS and antibody dosing required knowledge of group allocation, but outcome assessment and quantitative analysis were performed blinded to group wherever possible.

B16-F10 cells were detached using 0.25% trypsin-EDTA, washed and resuspended in sterile PBS. Mice were injected subcutaneously with 4 × 10^5^ B16-F10 cells on day 0. Bif-210 BEVs diluted in PBS (100 µL) or PBS alone were administered intravenously via the tail vein when tumours were palpable (day 6), then every 3 days for a total of three administrations (days 6, 9 and 12). Unless otherwise stated, the BEV dose was 10^9^ particles/mouse. Tumour growth was monitored by digital calliper measurement, and tumour volume was calculated as width^2^ × length × 0.52. Animals were euthanised on day 14 by gradual exposure to CO_2_. Core body temperature was measured from the abdominal region using an infrared thermometer.

### *In vivo* Ly6G depletion

To deplete Ly6G+ cells, mice received 200 µg/mouse anti-mouse Ly6G antibody (clone 1A8; 2B Scientific, BE0075-1-25MG) or rat IgG2a isotype control (anti-trinitrophenol; 2B Scientific, BE0089-25MG) by intraperitoneal injection (100 µL) every 2 days, commencing on day 5 post-B16-F10 implantation.

### Bone marrow isolation and BEV incubations

Femurs and tibias were removed under sterile conditions and marrow compartments flushed with sterile PBS using a 23G needle. Cells were centrifuged (300 g, 5 min), resuspended in ice-cold sterile 0.2% NaCl for 30 s to lyse red blood cells, and osmotic balance restored by addition of an equal volume of 1.6% ice-cold NaCl. Cells were resuspended in RPMI + 1% FBS for flow cytometry or plate-based assays. For *ex vivo* BEV incubations, bone marrow cells were seeded at 1 × 10^6^ cells/well in sterile 96-well non-tissue-culture-treated plates and incubated with PBS or BEVs (10^9^ particles/well unless otherwise stated) for 5 h. For extracellular DNA release assays, PMA (100 ng/mL) was used as a positive control for NETosis-associated extracellular DNA release ^37, 38^.

### Flow cytometry

Resected tumours were manually homogenised and samples prepared for flow cytometry as previously described ^45^. Blood was collected by cardiac puncture using 23G EDTA-coated needles immediately after euthanasia. Red blood cell lysis was performed before staining. Compensation was performed using AbC Total Antibody Compensation Beads (Thermo, A10497) and single-stained controls. Fluorescence minus one controls were used for gating, with gating strategies shown in Table 2. Samples were acquired on a BD LSRFortessa and analysed using FlowJo v10.9.0.

**Table 2:**
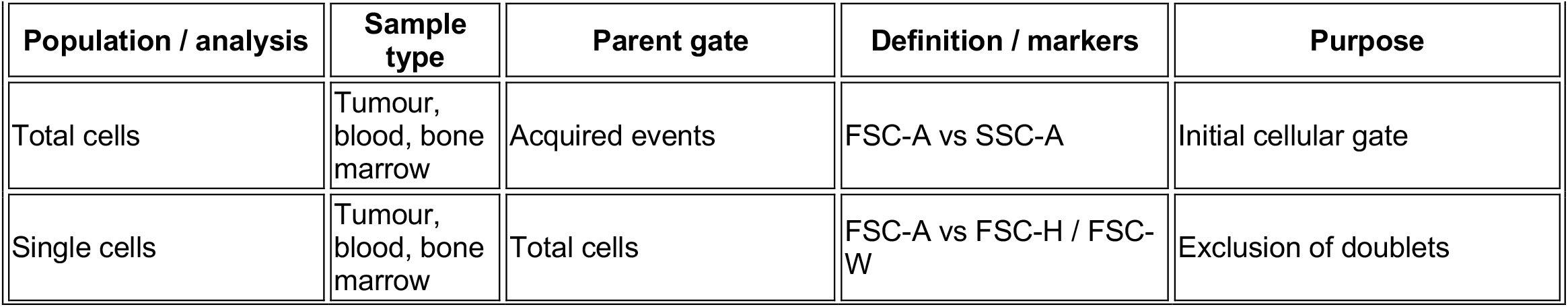
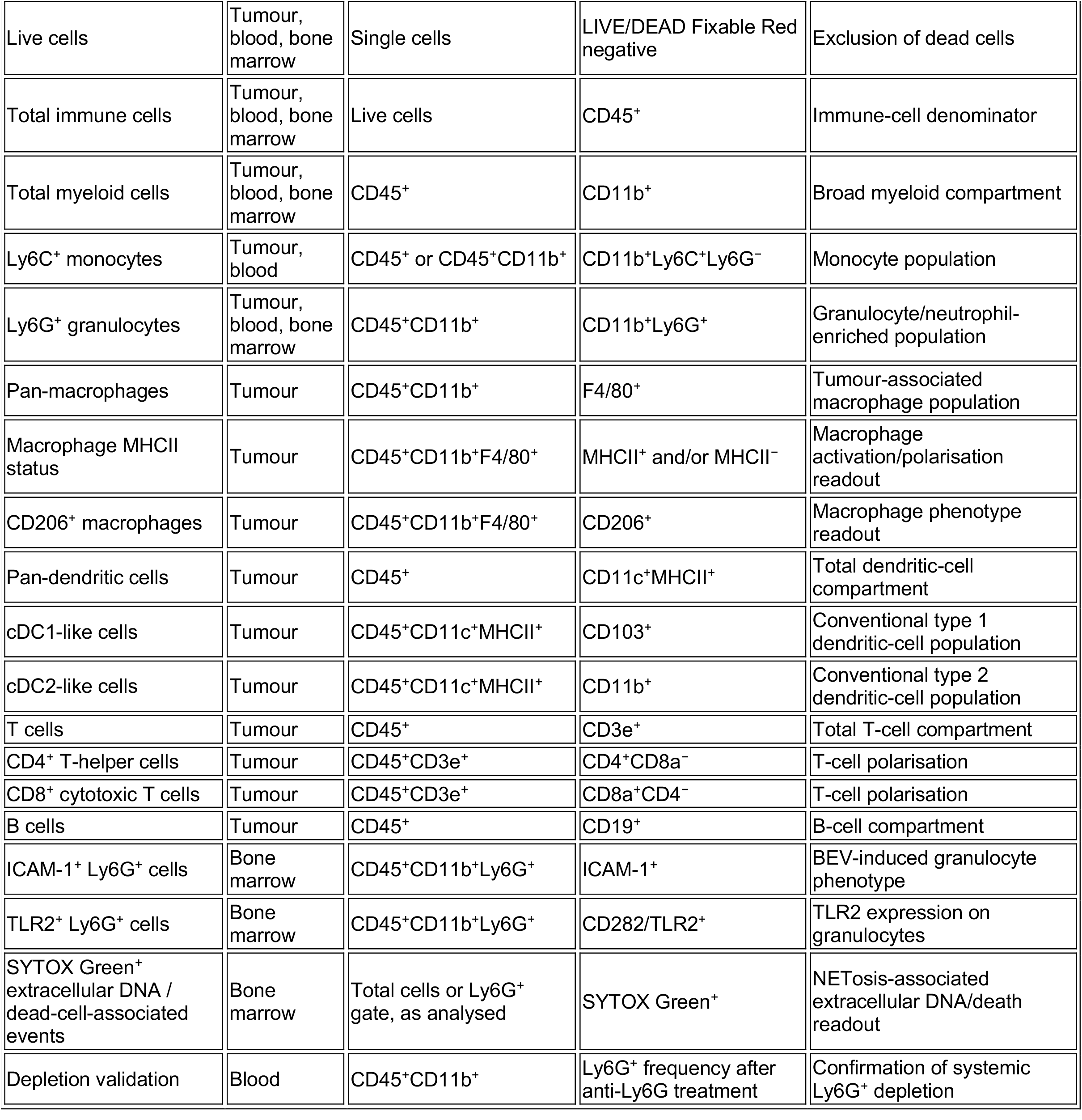
flow-cytometry gating strategies.

Antibodies/reagents: LIVE/DEAD Fixable Red (Thermo, L34971); SYTOX Green (Thermo, S7020); CD45-PerCP-Cy5.5 (30-F11, eBioscience, 45-0451-82); CD3e-APC (145-2C11, Thermo, 17-0031-81); CD4-FITC (RM4-4, eBioscience, 11-0043-81); CD8a-APC-Cy7 (53-6.7, Thermo, A15386); CD19-BV650 (6D5, BioLegend, 115541); CD11b-BV605 (M1/70, BD, 563015); Ly6C-eFluor450 (HK1.4, Invitrogen, 48-5932-82); Ly6G-APC-Cy7 (1A8, BD, 560600); ICAM-1-PE (YN1/1.7.4, BioLegend, 116107); F4/80-AF700 (BM8, BioLegend, 123129); F4/80-APC (BM8, Invitrogen, 17-4801-82); MHCII-PE-Cy7 (M5/114.15.2, Invitrogen, 25-5321-82); CD206-FITC (C068C2, BioLegend, 141704); CD11c-BUV395 (HL3, BD, 564080); CD103-BV711 (2E7, BioLegend, 121435); CD282/TLR2-PE-Cy7 (QA16A01, BioLegend, 153011).

### TLR reporter assays

HEK-Blue hTLR2 and HEK-Blue hTLR4 cells (InvivoGen) were cultured in RPMI-1640 supplemented with 10% FBS, 1% P/S, Normocin (100 µg/mL) and HEK-Blue selection cocktail. Cells were seeded at 5 × 10^4^ cells/well in 96-well plates containing HEK-Blue detection medium and treated with PBS, BEVs or positive controls (LTA-BS for TLR2 and LPS for TLR4; InvivoGen). After overnight incubation at 37°C, absorbance at 650 nm was measured using a microplate reader (BioTek, USA).

### MTS assay

B16-F10 cells were seeded at 2,000 cells/100 µL/well in flat-bottom tissue-culture-treated 96-well plates for 24 h, then media were replaced with fresh media containing 1% PBS or BEVs (10^9^ particles) for 24 h. Bone marrow cells were seeded at 1 × 10^6^ cells/100 µL/well in RPMI + 1% FBS and incubated with PBS or BEVs (10^9^ particles) for 5 h. MTS reagent (20 µL/well) was added for 1 h and absorbance was measured at 492 nm. Media-only values were subtracted, and viability/metabolic activity was presented relative to PBS-treated controls.

### Myeloperoxidase activity assay

MPO activity was assessed in tumour homogenates and whole blood. Tumour tissue (20 mg) from BEV- and PBS-treated animals was homogenised for 1 min using pellet pestles attached to a cordless motor. Mouse blood (200 µL) was incubated with BEVs (10^9^ particles) or PBS for 2 h. MPO activity was quantified using a colorimetric kit (Abcam, ab105136) according to the manufacturer’s instructions, and absorbance was measured at 412 nm.

### Cytokine quantification

THP-1 macrophages and THP1-Blue reporter monocytes were seeded at 1 × 10^5^ cells/well, and splenocytes at 1 × 10^6^ cells/well, in 96-well plates. Cells were treated with 10% BEVs (10^10^ particles/mL), 10% PBS or 10% LPS (200 ng/mL, Invitrogen) for 24 h. Supernatants were collected and stored at -80°C. Human IL-8 and mouse KC/CXCL1 were quantified using a customised U-PLEX MSD platform (MSD Inc.) according to the manufacturer’s instructions. Plates were read using a MESO QuickPlex SQ 120, and concentrations were calculated using MSD Discovery Workbench 4.0.

### Histopathological assessment

Lung, liver, spleen and kidney tissues were fixed overnight in 4% PFA at 4°C, transferred to 100% ethanol, processed using a Leica Tissue Processor ASP300S and embedded in paraffin wax using an EG1150H embedding station (Leica Biosystems). FFPE sections (4 µm) were stained with haematoxylin and eosin using an automated multistainer (Leica Biosystems, ST5020) according to the manufacturer’s instructions.

### Quantification and statistical analysis

Graphs and statistical analyses were generated using GraphPad Prism 10. Unless otherwise stated, two-group comparisons used two-tailed Student’s t-tests; multi-group comparisons used one- or two-way ANOVA with Tukey’s or Dunnett’s multiple-comparison tests as appropriate; non-parametric data were analysed using Mann-Whitney tests. Data were inspected for distribution and variance where appropriate; if assumptions for parametric analysis were not met, non-parametric alternatives were used. Data are presented as mean ± SEM unless otherwise stated. Significance thresholds were: p > 0.05, not significant; *p < 0.05; **p < 0.01; ***p < 0.001; ****p < 0.0001. Exact n values and statistical tests are provided in figure legends. Effect sizes with confidence intervals are not currently reported; group differences are presented using individual data points, mean ± SEM, graph-indicated significance levels and n values.

## RESOURCE AVAILABILITY

### Lead contact

Further information and requests for resources should be directed to the lead contact, Stephen D. Robinson.

### Materials availability

Bif-210 and BEV preparations generated in this study may be available from the lead contact subject to institutional approvals and material transfer agreements.

### Data and code availability

Raw data supporting the conclusions of this study will be made available by the authors without undue reservation. This study did not generate custom code.

## AUTHOR CONTRIBUTIONS

Conceptualisation: ADN, AJ, CAP, LJH, SDR. Methodology: ADN, AJ, CAP, MR, NI, LM, RS, SDR. Formal analysis: ADN, AJ, MR, SDR. Investigation: ADN, AJ, CAP, MR, NI, LM. Resources: SRC, LJH, RS, SDR. Training and specialist BEV support: RS. Writing - original draft: ADN, SDR. Writing - review and editing: ADN, AJ, CAP, MR, NI, LM, RS, SRC, LJH, SDR. Visualisation: ADN, AJ, LJH, SDR. Supervision: SRC, LJH, SDR. Funding acquisition: SRC, LJH, SDR.

## ACKNOWLEDGMENTS

SRC, SDR and RS acknowledge support from the Biotechnology and Biological Sciences Research Council (BBSRC). This research was supported by the BBSRC Institute Strategic Programme Grants Gut Microbes and Health (BB/R012490/1) and its constituent projects (BBS/E/F/000PR10353, BBS/E/F/000PR10355 and BBS/E/F/000PR10356), and Food Microbiome and Health (BB/X011054/1) and its constituent projects (BBS/E/QU/230001B and BBS/E/QU/230001C). This work was supported by the Wellcome Trust-funded EDESIA: Plants, Food and Health PhD programme [218467/Z/19/Z]. AJ was supported by the BBSRC Norwich Research Park Bioscience Doctoral Training Partnership grant BB/M011216/1 (supervisors L.J.H. and S.R.C.; student A.J.). We thank the Quadram Institute Bioinformatics Core and the Quadram Institute Bioimaging Facility at Quadram Institute Bioscience for expert support with data analysis and imaging.

## FIGURE LEGENDS

**Supplementary Figure 1.**
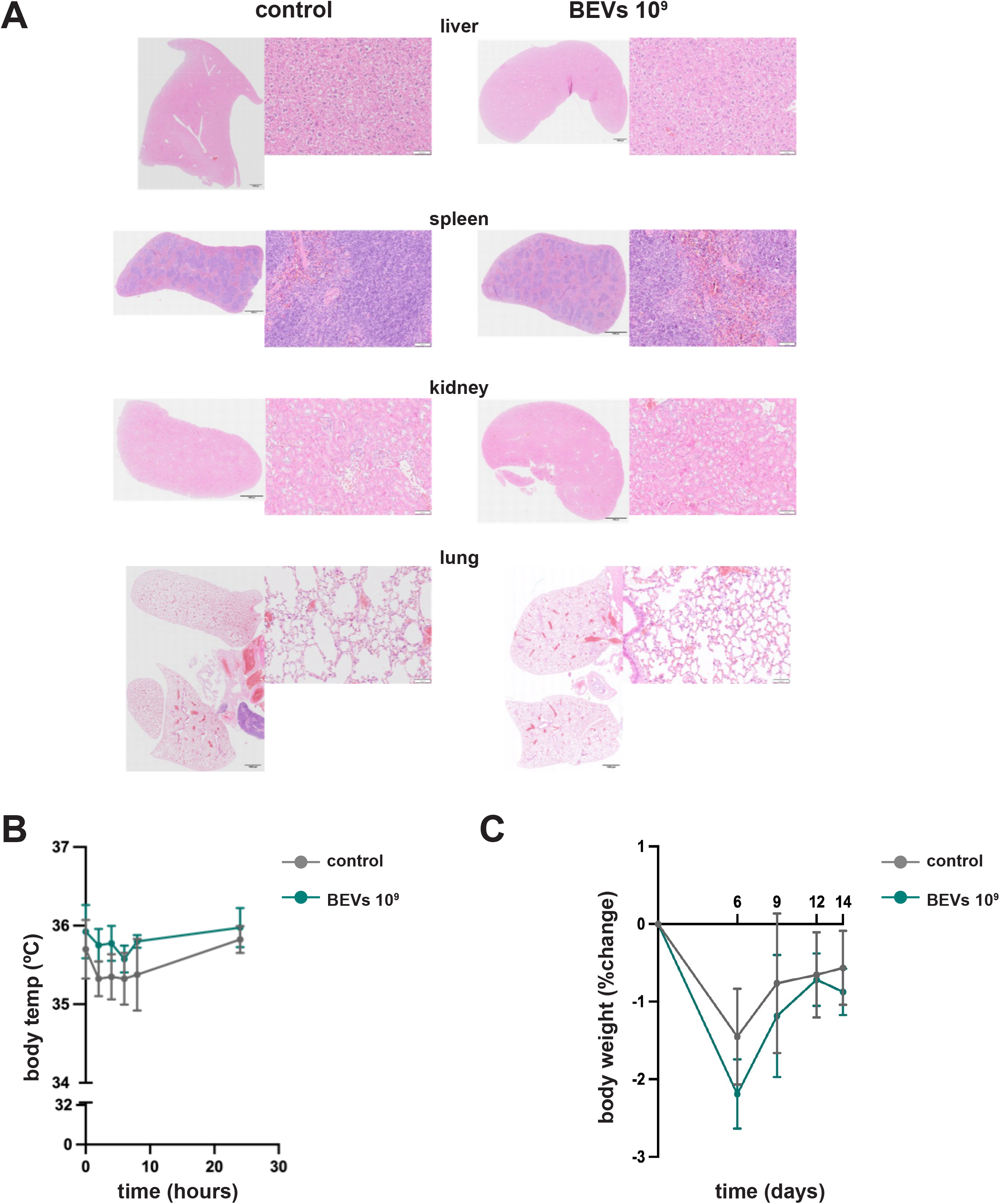
Repeated intravenous Bif-210 BEV administration shows no overt toxicity in mice. Non-tumour-bearing C57BL/6 mice received three intravenous doses of Bif-210 BEVs (10^9^ particles/mouse) or PBS using the same dosing schedule as tumour-bearing mice. (A) Representative H&E-stained sections of liver, spleen, kidney and lung from PBS- and BEV-treated mice. Scale bars = 1,000 µm for whole-organ images and 50 µm for higher-magnification images. (B) Core body temperature monitored from baseline every 2 h until 8 h and again at 24 h after administration. (C) Body weight monitored over the 14-day dosing period and expressed as percentage change from baseline. Data are mean ± SEM; statistical tests as described in Methods.

**Supplementary Figure S2.**
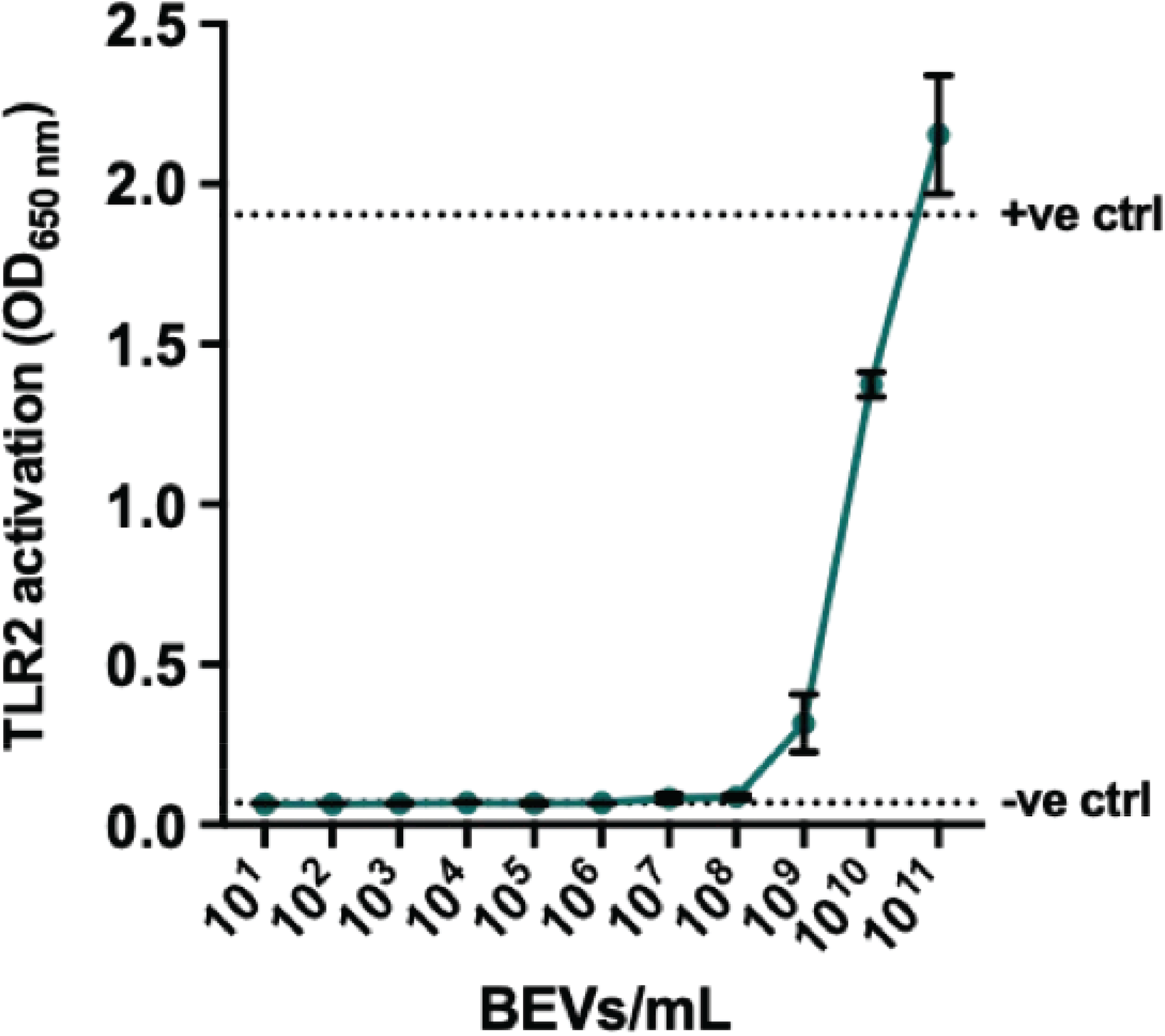
Bif-210 BEVs activate TLR2 in a dose-dependent manner. HEK-Blue hTLR2 reporter cells were stimulated with increasing concentrations of Bif-210 BEVs. TLR2 activation was measured by absorbance at 650 nm. Data are shown as mean ± SEM. Dotted lines indicate the positive and negative controls.

